# Towards a new history and geography of human genes informed by ancient DNA

**DOI:** 10.1101/003517

**Authors:** Joseph K. Pickrell, David Reich

**Affiliations:** New York Genome Center, New York, NY; Department of Biological Sciences, Columbia University, New York, NY; Department of Genetics, Harvard Medical School, Boston, MA; Howard Hughes Medical Institute, Harvard Medical School, Boston, MA; Broad Institute of MIT and Harvard, Cambridge, MA

## Abstract

Genetic information contains a record of the history of our species, and technological advances have transformed our ability to access this record. Many studies have used genome-wide data from populations today to learn about the peopling of the globe and subsequent adaptation to local conditions. Implicit in this research is the assumption that the geographic locations of people today are informative about the geographic locations of their ancestors in the distant past. However, it is now clear that long-range migration, admixture and population replacement have been the rule rather than the exception in human history. In light of this, we argue that it is time to critically re-evaluate current views of the peopling of the globe and the importance of natural selection in determining the geographic distribution of phenotypes. We specifically highlight the transformative potential of ancient DNA. By accessing the genetic make-up of populations living at archaeologically-known times and places, ancient DNA makes it possible to directly track migrations and responses to natural selection.

## Introduction

Within the past 100,000 years, anatomically modern humans have expanded to occupy every habitable area of the globe. The history of this expansion has been explored with tools from a number of disciplines, including linguistics, archaeology, physical anthropology, and genetics. How did we get to where we are today?

Genetic information, since it is passed from generation to generation so faithfully, contains powerful information about population history. Technological advances have now made all of this information easily accessible. The great synthesis of genetic data with historical, archaeological and linguistic information, “The History and Geography of Human Genes” (Cavalli-Sforza et al., 1994), was written based on data from around one hundred protein polymorphisms. However, it is now possible to genotype millions of polymorphisms in thousands of individuals using high-throughput sequencing at an affordable cost (Abecasis et al., 2010).

With the advent of extensive genome-wide data, geneticists have been able to propose more refined models for the history of the human expansion across the globe. Probably the most popular model of this type, the “serial founder effect” model, is one in which humans expanded around the world in a series of successive population splits and range expansions into unoccupied territory, with little subsequent migration. Under this model, the present-day inhabitants of any region are largely descended from the first inhabitants of the region (Harpending and Eller, 2000; Harpending and Rogers, 2000; Hellenthal et al., 2008; Henn et al., 2012a; Prugnolle et al., 2005; Ramachandran et al., 2005).

It is now clear that such models are inaccurate for many populations. The present-day inhabitants of many regions of the world do not descend in simple ways from the populations that lived in the same locations in the distant past. Instead, human populations have periodically experienced major demographic upheavals, such that much of the geographic information about the first human migrations has been overwritten by subsequent population admixture or replacement. This idea has a long (albeit controversial) history in archaeology; it has been debated for decades whether sudden changes in material culture apparent in the archaeological record can be attributed to population movements (Renfrew and Bahn, 1996). In genetics, the series of discoveries documenting the importance of mixture and migration in human history have come in part from analysis of populations living today (e.g. Hellenthal et al., 2014; Loh et al., 2013; Patterson et al., 2012), but also increasingly from genetic analysis of ancient human remains (e.g. Raghavan et al., 2013; Skoglund et al., 2012).

In this paper, we argue that these results motivate a systematic re-evaluation of human history using modern genomic tools—a new “History and Geography of Human Genes” that exploits many orders of magnitude more data than the original synthesis—and we emphasize the potential for using ancient DNA to answer outstanding questions. Other recent reviews have had different emphases, or have covered methodological approaches in more detail than we do here (Alves et al., 2012; Colonna et al., 2011; Novembre and Ramachandran, 2011; Pinhasi et al., 2012; Stoneking and Krause, 2011; Veeramah and Hammer, 2014; Wall and Slatkin, 2012)

## Re-evaluation of the “serial founder effect” model

For about the last ten years, genetic studies of early human history have often been framed around the “serial founder effect” model. This model, initially proposed by Harpending, Eller, and Rogers (Harpending and Eller, 2000; Harpending and Rogers, 2000), gained popularity with the publication of two papers (Prugnolle et al., 2005; Ramachandran et al., 2005) that observed that heterozygosity (the average number of differences between two random copies of a genome in a population) declines approximately linearly with geographic distance from Africa. This pattern, initially identified in genome-wide microsatellite genotypes from 53 worldwide human populations, was subsequently confirmed in large single nucleotide polymorphism (SNP) datasets (Li et al., 2008).

The observation of declining human genetic variation with increasing geographic distance from Africa was interpreted as being consistent with a “serial founder effect” model for human dispersal. In this type of model (Figure 1A), the peopling of the globe proceeded by an iterative process in which small bands of individuals pushed into unoccupied territory, experienced population expansions, and subsequently gave rise to new small bands of individuals who then pushed further into unoccupied territory. This model has two important features: a large number of expansions into new territory by small groups of individuals (and concurrent bottlenecks), and little subsequent migration. As Prugnolle et al. (2005) wrote: “what is clear…is that [this] pattern of constant loss of genetic diversity along colonisation routes could only have arisen through successive bottlenecks of small amplitude as the range of our species increased… The pattern we observe also suggests that subsequent migration was limited or at least very localized”.

**Figure 1.**
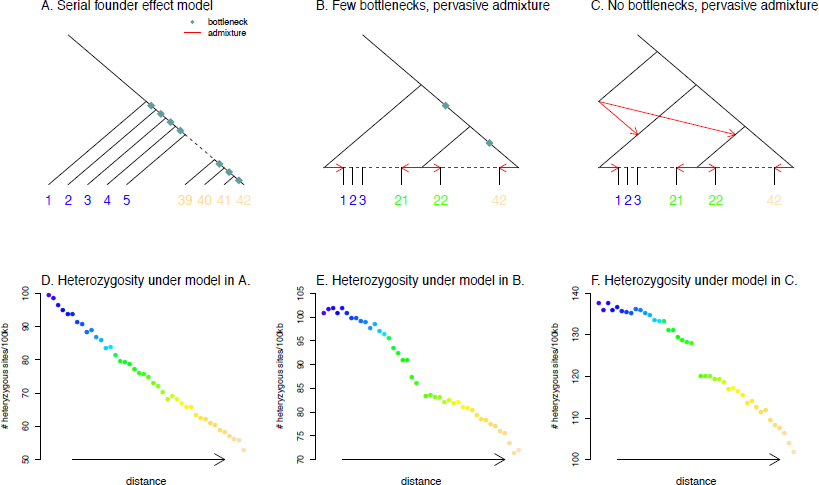
A negative correlation between heterozygosity and geographic distance from a source population can be generated by qualitatively different, historically plausible demographic models. We simulated genetic data under different demographic models and calculated the average heterozygosity in each simulated population. **A.** Schematic of a serial founder effect model. **B.** Schematic of a demographic model with two bottlenecks and extensive admixture. **C.** Schematic of a demographic model with no bottlenecks and extensive admixture. **D,E,F**. Average heterozygosity in each population simulated under the demographic models in A,B, and C respectively. Each point represents a population, ordered along the x-axis according as in A.

This model has been influential in many fields. If the serial founder event model is correct, then the difficult problem of identifying the geographic origin of all modern humans is reduced to the simpler problem of finding the geographic region where people have the most genetic diversity (Ramachandran et al., 2005). This idea has been used extensively in discussions of human origins (e.g. Henn et al., 2011; Schlebusch et al., 2012). Outside of genetics, the serial founder effect model has been used as a framework to interpret data from linguistics (Atkinson, 2011), physical anthropology (Betti et al., 2012; von Cramon-Taubadel and Lycett, 2008; Hanihara, 2008; Manica et al., 2007), material culture (Rogers et al., 2009), and economics (Ashraf and Galor, 2013).

A caveat of the serial founder effect model is that many other historical models can produce qualitatively similar patterns of genetic diversity (e.g. Amos and Hoffman, 2010; DeGiorgio et al., 2009). Producing this empirical pattern simply requires that the average time to the most recent common ancestor between two chromosomes in a population depend on the distance of that population from Africa (DeGiorgio et al., 2009). To illustrate this point, we simulated genetic data using *ms* (Hudson, 2002) under three demographic models: the serial founder effect model (Figure 1A), a model with two large founder effects and extensive subsequent population mixture (Figure 1B), and a model without bottlenecks but with archaic admixture (from an anciently diverged population like Neanderthals) as well as more extensive recent population mixture (Figure 1C). Details of the simulation parameters are in the Supplementary Information. We simulated 1,000 regions of 100kb under each model, and plotted the average heterozygosity in each population. In all three cases (Figure 1D,E,F) we recapitulate an approximately linear decline in heterozygosity with distance from a source. These simulations show that the main observation that has been marshaled in support of the serial founder effect model is also consistent with very different histories. Other types of evidence are necessary to distinguish between these very distinct and in our view equally plausible models.

## The current inhabitants of a region are often poor representatives of the historical populations who lived in the same locations

A parsimonious assumption underlying much work on human population history is that populations living in a geographic region today are largely descended from the first anatomically modern humans to arrive in the region. Recent work, however, has shown that the truth is far from parsimonious. Instead, long-range migration and concomitant population replacement or admixture has occurred often enough in human history that the present-day inhabitants of many places in the world often bear little genetic relationship to the more ancient peoples of the same region.

The present-day populations of the Americas represent a recent example of this phenomenon. It is well known that the populations of the Americas experienced demographic upheaval after the arrival of Europeans and Africans around 500 years ago. Accounting for mixture between Europeans, Africans and Native Americans is crucial to understanding population structure in the Americas. In fact, most of the ancestry of present-day populations in the Americas is not derived from the Native Americans who were the sole inhabitants of the region less than half a millennium ago (Bryc et al., 2010; Gravel et al., 2013; Johnson et al., 2011; Moreno-Estrada et al., 2013; Price et al., 2007; Wang et al., 2007). Another recent example is Australia, where accounting for massive European migration over the last couple of hundred years is crucial for understanding population structure (McEvoy et al., 2010).

Further back in time, the present-day hunter-gatherer and pastoralist populations of Siberia are often treated as surrogates for the populations that first crossed the Bering land bridge to people the Americas beginning more than 15,000 years ago (e.g. Lell et al., 2002; Santos et al., 1999; Starikovskaya et al., 2005; Wang et al., 2007). However, just in the last year, genetic data showed that this assumption is wrong. The sequencing of DNA from an individual who lived ~24,000 years before the present in the Lake Baikal region of Siberia, and a ~17,000 year old individual from the same region, showed that these ancient individuals (from time points prior to the time the Americas were thought to be peopled) are phylogenetically more closely related to present-day Native Americans than are present-day Siberians (Raghavan et al., 2013). These ancient individuals appear to have been members of an “Ancient North Eurasian” population that no longer exists in unmixed form, but that admixed substantially with the ancestors of both present-day Europeans and Native Americans (Patterson et al., 2012; Raghavan et al., 2013; Lazaridis et al., 2013). In contrast, most of the present-day indigenous populations of Siberia are more closely related to East Asians, indicating that the present-day indigenous population of Siberia is descended in large part from populations that arrived in the region after the end of the last ice age (Raghavan et al., 2013).

Ancient DNA studies have also revealed population turnover over shorter timescales (Bramanti et al., 2009; Brandt et al., 2013; Haak et al., 2005, 2010; Keller et al., 2012; Lazaridis et al., 2013; Malmström et al., 2009; Skoglund et al., 2012). A major debate in the last few decades among archaeologists and geneticists has been about whether the arrival of agriculture in Europe involved the spread of people or culture. Skoglund et al. (2012) addressed this controversy by analyzing genome-wide genetic data from Swedish hunter-gatherer and agriculturalist populations that lived around 5,000 years ago. The farmer population appears most genetically similar to southern Europeans today, while the hunter-gatherers are more similar to northern Europeans. Thus, at least in Scandinavia, the spread of agriculture was accompanied by the spread of people. Mitochondrial DNA studies have shown that such population turnovers accompanied the arrival of agriculture throughout Europe (Bramanti et al., 2009; Brandt et al., 2013; Haak et al., 2005, 2010).

The arrival of farmers was not the end of pre-historic population turnover in Europe (and we do not even discuss here the turnovers that occurred in the few thousand years in Europe since the invention of writing; see Davies, 1998; Hellenthal et al., 2014; Moorjani et al., 2011; Ralph and Coop, 2013). The most notable study in this regard is the one of Brandt et al. (2013), which assembled mtDNA data from 364 human samples from archaeological cultures ranging from the early Neolithic to the Bronze Age from the same geographic area in Germany. This study documented discontinuity between people of early and late Neolithic cultures, with people of late Neolithic cultures bearing more relatedness to the present-day populations of Eastern Europe and Russia than do people of early Neolithic cultures. Thus, major demographic turnover has happened at least twice over the course of the last eight thousand years of European prehistory. This makes inferences about the inhabitants of Europe *tens* of thousands of years ago based on the locations of people today unreliable.

Evidence of major demographic changes in the last several thousand years has also accumulated in parts of the world where no ancient DNA is (yet) available:

- In India, nearly all people today are admixed between two distinct genetic groups, one most closely related to present-day Europeans, Central Asians, and Near Easterners, and one most closely related to an isolated population living in the Andaman islands (Reich et al., 2009). Moorjani et al., 2013 showed that much of this admixture occurred within the last 4,000 years.
- In North Africa, nearly all people today descend from admixture between populations related to those present today in western Africa and in the Near East (Henn et al., 2012b; Rosenberg et al., 2002; Tishkoff et al., 2009). Some of this admixture can be dated to within the last few thousand years (Hellenthal et al., 2014), indicating that much of the ancestry from these populations does not descend continuously from the Stone Age peoples of North Africa.
- In sub-Saharan Africa, genetic studies have documented multiple examples of populations with ancestry from disparate sources. Populations that speak languages with heavy use of click consonants in eastern Africa (Hadza and Sandawe) result from admixture involving a population related to hunter-gatherer populations southern Africa (Khoisan) (Pickrell et al., 2012), suggesting that Khoisan-related populations were once more widespread than they are today. Further, populations in eastern and southern Africa have been influenced by gene flow from west Eurasian populations in the last 3,000 years (Pagani et al., 2012; Pickrell et al., 2014). And in Madagascar, all populations derive approximately half of their ancestry from Austronesian speaking migrants from southeast Asia (Pierron et al., 2014).

Beyond case studies, Patterson et al. (2012), Loh et al. (2013), and Hellenthal et al. (2014) applied tests for admixture to diverse sets of population from around the globe and found that nearly all populations show evidence of admixture. To illustrate the degree to which this admixture involved populations that today are geographically distant, in Figure 2 we present an analysis based on combining data for 103 worldwide populations from a number of sources (Altshuler et al., 2010; Behar et al., 2010; Henn et al., 2011; Li et al., 2008; Pagani et al., 2012; Schlebusch et al., 2012) and running a simple three-population test for admixture (Reich et al., 2009) on all populations (see Supplementary information for details). For all populations with significant evidence of admixture, we identified the present-day populations that are most closely related to the admixing populations.

**Figure 2.**
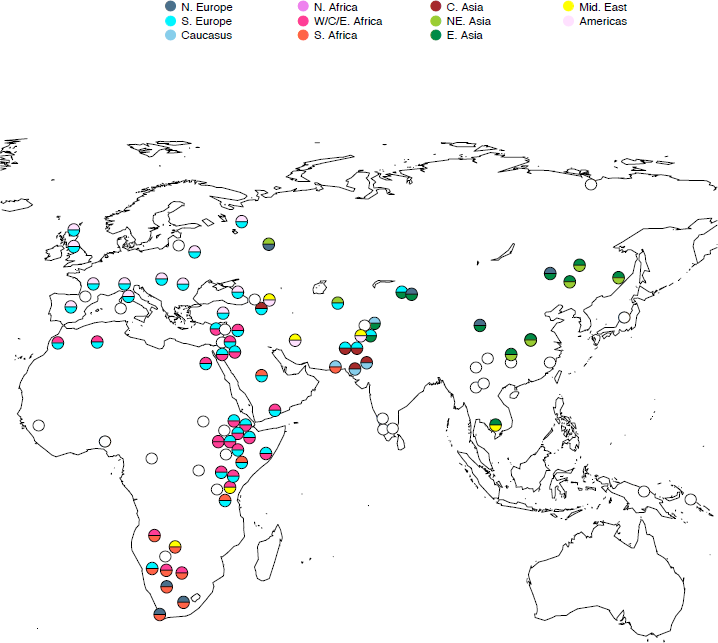
A birds-eye view of admixture in human populations. We performed a three-population test for admixture (Reich et al., 2009) on 103 worldwide populations. Circles show the approximate current geographic locations of all tested populations (except for populations in the Americas, which are not plotted for ease of display). Filled circles represent populations identified as admixed, and the colors represent the current geographic labels of the inferred admixing populations. Empty circles represent populations with no statistically-significant evidence for admixture in this test.

The results are visually startling. Figure 2 shows the geographic locations of all the populations, along with the locations of the best present-day proxies of their ancestral populations. Admixture between populations related to ones that are now geographically distant is evident in most populations of the world. For example, Native American-related ancestry is present throughout Europe (Patterson et al., 2012), likely reflecting the genetic input from the Ancient Northern Eurasian population related to Upper Paleolithic Siberians (Lazaridis et al., 2013; Patterson et al., 2012) both into the Americas (most likely prior to 15,000 years ago) and into Europe (Lazaridis et al., 2013; Raghavan et al., 2013). Also, Near Eastern-related ancestry is found in Cambodia (Pickrell and Pritchard, 2012), likely due to mixture from an ancestral South Asian population that was itself an admixed population containing ancestry related to present-day Near Easterners (Hellenthal et al., 2014). The test we use as the basis for Figure 2 detects only the largest signals of admixture in the data, and cannot detect complete population replacement. The true population history is likely to have been even more complex.

These examples show that the parsimonious assumption often made in population genetics—that the populations in a given region today are largely descended from the inhabitants of that geographic region in the distant past—is rarely true. Combined with the simulations presented in Figure 1, these results demonstrate that the serial founder effect model is no longer a reasonable null hypothesis for modeling the ancient spread of anatomically modern humans around the globe. This motivates a search for alternative models that may better fit the data.

The type of model that is emerging from the data is presented in Figure 3. In this model, the major land masses of the world have all been peopled multiple times, with population replacement and/or admixture happening periodically throughout history. As new data become available, this model will likely be refined and completely re-written in parts.

**Figure 3.**
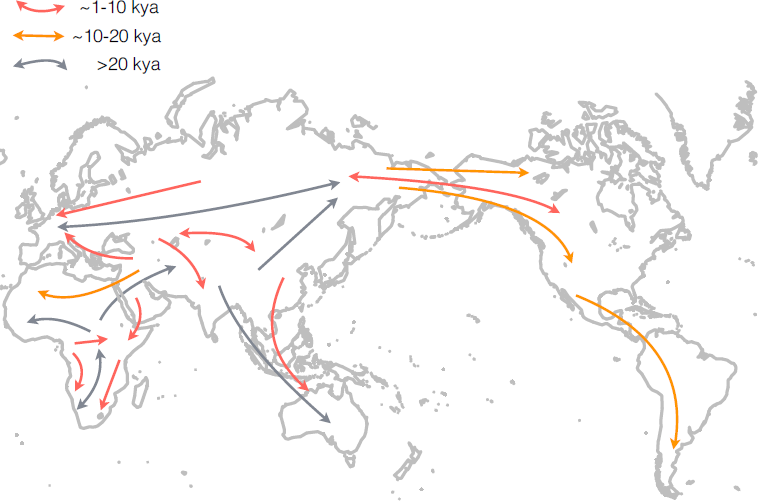
A rough guide to genetically documented population movements in the history of anatomically modern humans. Arrows represent population movements, and the color of each arrow represents the approximate time of the migration.

## Ancient DNA: a transformative source of information about the past

The types of models that are most useful for making sense of human history are ones that specify geographic as well as temporal information; that is, those that make statements of the form “a population from location X moved to location Y during time period Z”. For example, one might wish to test whether the first pastoralists in southern Africa arrived as migrants from eastern Africa after around 2,500 years ago (the time when evidence of pastoralism in southern Africa appears in the archaeological record), against the alternative hypothesis that a pastoralist lifestyle was adopted by indigenous people who learned it through cultural transmission. Testing this hypothesis using DNA from individuals today involves assuming that populations in southern and eastern Africa today are representative of the populations in southern and eastern Africa at the times of interest. As discussed above, this assumption is often untrue.

Studies of DNA from ancient human remains offer a way around this limitation. Rather than studying the past by the traces it has left in present-day people—which is problematic since human individuals and even whole populations are capable of migrating hundreds or thousands of kilometers in a lifetime—ancient DNA offers the ability to analyze the genetic patterns that existed at a particular time at a geographical location. This allows direct inference about the relationships of historical populations to each other and to populations living today. Below we list some surprising findings about human history based on ancient DNA, which could not have been obtained without this source of information:

- There was archaic Neanderthal gene flow into the ancestors of all anatomically modern humans outside African 37,000-85,000 years ago (Sankararaman et al., 2012). This gene flow contributed on the order of 2% of the genetic ancestry of non-Africans (Green et al., 2010; Prüfer et al., 2014). This finding was suggested by genetic analysis prior to the use of ancient DNA to document this admixture event (Plagnol and Wall, 2006). However, the consensus about whether admixture occurred only changed after ancient DNA evidence showed that the deeply diverged segments in present-day non-Africans are related to Neanderthals (Green et al., 2010).
- A previously unknown archaic population that was neither Neanderthal nor modern human was present in Siberia before 50,000 years ago (Krause et al., 2010; Reich et al., 2010). The discovery of the “Denisovans” shows how ancient DNA can reveal “genomes in search of a fossil”; populations that were not expected at all based on the archaeological or fossil record.
- There was gene flow from a population related to the Denisovans into the ancestors of present-day aboriginal people from New Guinea, Australia, and the Philippines (Cooper and Stringer, 2013; Meyer et al., 2012; Prüfer et al., 2014; Reich et al., 2010, 2011). It is notable that the populations today that contain the largest proportion of Denisovan ancestry are in Oceania, far from where the Denisovan bone was found in Siberia, a finding that again would not have been expected in the absence of ancient DNA data.
- There was admixture between an Ancestral North Eurasian population and a population related to present-day East Asians in the ancestry of Native Americans prior to the diversification of Native American populations in the New World (Raghavan et al., 2013).
- There were at least two important changes in population structure in central Europe over the course of the last 9,000 years (Bramanti et al., 2009; Brandt et al., 2013; Haak et al., 2005, 2010; Keller et al., 2012; Lazaridis et al., 2013; Malmström et al., 2009; Skoglund et al., 2012).
- Populations in the Americas have experienced multiple episodes of population admixture or turnover, such that an individual living in Greenland around 4,000 years ago is more closely related to populations currently in Siberia than to present-day Greenland Inuit populations (Rasmussen et al., 2010), and an individual living in North America around 12,000 years ago is more closely related to present-day Native South American populations than to Native North American populations (Rasmussen et al., 2014).

Ancient DNA results are so regularly surprising that almost any measurement is interesting: new historical discoveries have been made in virtually every ancient DNA study that has been carried out. The reason why ancient DNA studies are so informative is that the technology provides a tool to measure quantities that were previously unmeasurable. In this sense, the value of ancient DNA technology as a window into ancient migrations is analogous to the 17^th^ century invention of the light microscope as a window into the world of microbes and cells.

## Scientific opportunities for ancient DNA studies

Moving forward, ancient DNA studies afford major opportunities in two areas: studies of population history, and studies of natural selection.

The literature on ancient mitochondrial DNA contains a number of promising study designs. For example, one might sample multiple individuals from different archaeological cultures at a single time point (Malmström et al., 2009). Such a “horizontal time slice” allows a snapshot of population structure over a broad geographic region, which can then be compared to the relatively complete picture of population structure today. An alternative study design (like that used by Brandt et al., 2013) is to take a single geographic location and sample individuals from multiple time points. This “vertical time slice” allows direct quantification of changes in population composition over time.

Mitochondrial DNA studies have major drawbacks compared with analysis of the whole genome, however (Ballard and Whitlock, 2004). First, mtDNA is inherited maternally, and thus does not capture any information about the history of males (which may differ from that of females due to sex-biased demographic processes). More importantly, a study of a single locus (or two loci if the Y chromosome is included to avoid the first limitation) has less statistical resolution for studies of history than do studies of the nuclear genome. The reason for this is that whole genome studies of an individual obtain information about hundreds of thousands of that individual’s ancestors, not just those on a single lineage. It is thus important that the more advanced study designs of the mtDNA studies be combined with analysis of the more informative autosomal DNA.

What outstanding questions in population history can be addressed with these study designs? One important question is whether changes in populations over time are typically gradual—due to consistent, low-level gene flow between neighboring populations—or punctate, with migration events rapidly altering the population genetic composition of a region. One line of work on modeling human history explicitly assumes the latter (Hellenthal et al., 2014; Lipson et al., 2013; Loh et al., 2013; Patterson et al., 2012; Pickrell and Pritchard, 2012; Reich et al., 2009). If this assumption is unfounded, however, other models that accommodate continuous gene flow (e.g. Excoffier et al., 2013; Gutenkunst et al., 2009) may be more appropriate. In principle, distinguishing between these possibilities with a time series of ancient DNA is straightforward; in Figure 4A we show how the predictions differ.

**Figure 4.**
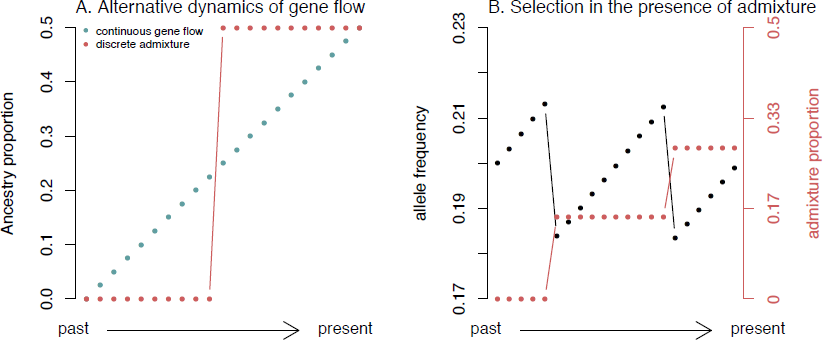
Time series allow tests of different models about human history. **A.** Different scenarios for the dynamics of gene flow. We show idealized time series of admixture proportions in a population under models of continuous gene flow and discrete admixture. **B.** Selection in the presence of admixture. We show idealized time series of the allele frequencies at a selected allele and admixture proportions in a population. Note that despite selection the allele does not show any net change in allele frequency.

Ancient DNA is also a promising way to address ongoing historical debates about the origins of different populations. Do linguistic isolates like the Basques in Europe have more ancestry from the pre-farming Mesolithic populations of the region than do their neighbors (Behar et al., 2012)? When did the west Eurasian-related population(s) that admixed with most Indian populations first appear in South Asia (Reich et al., 2009)? Did the first modern human inhabitants of East Asia descend from an earlier out-of-Africa migration than the populations living in East Asia today (Rasmussen et al., 2011; Reich et al., 2011)? Answering these questions will require a time series of snapshots of human genetic structure, combining the two types of study design mentioned above. If experience is a guide, this type of information will uncover additional unexpected aspects of human history.

The admixture and population replacements identified by ancient DNA also have implications for studies of natural selection. It is often assumed that populations have been in their current geographic locations long enough to adapt to their local conditions. This is explicit in approaches that look for correlations between environmental variables and allele frequencies (e.g. Coop et al., 2010; Hancock et al., 2010a; Huerta-Sánchez et al., 2013) and implicit in studies that interpret selected loci in terms of the current locations of populations (e.g. Simonson et al., 2010).

To what extent are the geographic distributions of selected alleles today indicative of the geographic distributions of selective pressures? (Note that the answer to this question may depend on the selective pressure in question). In individual cases, these two distributions are highly correlated; the classic example is the correlation between malaria incidence and disorders of hemoglobin (Kwiatkowski, 2005). For other cases, the correlation is imperfect. For example, alleles causing light skin pigmentation are at high frequency in northern Africa (Norton et al., 2007). If the selective pressure causing light skin is indeed a relative lack of ultraviolet radiation (Jablonski and Chaplin, 2000), it seems reasonable to expect that people living around the Sahara desert will be unaffected by this selection pressure. It is thus likely that the high frequency of alleles causing light skin pigmentation in north Africa was caused by the arrival of lightly-pigmented people (Henn et al., 2012b).

More generally, it has been observed that allele frequencies at loci under natural selection in humans tend to track neutral population structure rather than any obvious geographical variation in selection pressures (Coop et al., 2009; Granka et al., 2012). One factor contributing to this observation may be that selection pressures on individual loci are relatively weak due to the quantitative nature of phenotypes (Hancock et al., 2010b; Hernandez et al., 2011; Pritchard et al., 2010; Turchin et al., 2012). Another contributing factor may be that population movements over the last several thousand years have to some extent decoupled the geographic distributions of selected alleles from the geographic distributions of selective pressures. From the point of view of an individual allele, the movement of populations acts as a form of fluctuating selection; as populations move from environment to environment, the selection coefficient on an allele may change in both sign and magnitude (assuming a fixed selection coefficient in a given environment). This means that the environment that dominated the allele frequency trajectory may not be the environment that the allele is found in today.

Ancient DNA is also a potentially transformative tool for understanding human adaptation more generally. Nearly all methods for detecting positive selection at the genetic level are based around the principle that a selected allele changes frequency more quickly than a neutral locus. Tracking the trajectories of individuals over time allows direct access to this information, and makes it possible to infer precisely when in time (and with more certainty where geographically) genetic changes began to arise (Wilde et al., 2014). In fact, in the presence of admixture it is simple to create scenarios where the trajectory of the selected allele allows one to identify selection despite there being no net change in allele frequency (Figure 4B). To date, studies of selection over time have been limited either to small sample sizes (Lazaridis et al., 2013; Olalde et al., 2014) or small numbers of sites (Krüttli et al., 2014; Malmström et al., 2010; Plantinga et al., 2012; Wilde et al., 2014). However, whole genome technologies should make it possible to interrogate many thousands of phenotypically-relevant variants simultaneously.

## Taming the wild west of ancient DNA

Realizing the potential of ancient DNA studies will require systematic approaches. However, ancient DNA studies today are often more spectacular than systematic.

The paradigmatic approach to ancient DNA research in the era of high-throughput sequencing involves identifying a “golden” archaeological sample that yields usable DNA and obtaining a complete or partial genome sequence from it (e.g. Green et al., 2010; Keller et al., 2012; Meyer et al., 2012; Prüfer et al., 2014; Raghavan et al., 2013; Rasmussen et al., 2010). To some extent, the discovery of analyzable samples has been the main driver of the scientific questions asked. However, this opportunistic approach will not be driving ancient DNA research in a few years’ time. The future of ancient DNA research will involve a return to a more systematic approach: hypothesis-driven sampling across time and space, and analysis of much larger sample sizes.

Ancient DNA research has three major experimental challenges and two computational challenges that need to be jointly addressed by researchers who wish to access this transformative technology.

The first challenge in ancient DNA research is the danger of contamination from archaeologists or lab researchers who have handled the sample. This is a recurring problem (Cooper and Poinar, 2000; Green et al., 2006, 2009; Hedges and Schweitzer, 1995; Pääbo, 1985; Pääbo et al., 2004; Wall and Kim, 2007; Willerslev et al., 2004). While experimental guidelines (e.g. Gilbert et al., 2005; Hofreiter et al., 2001; Pääbo et al., 2004) as well as laboratory methods for reducing the possibility of contamination (Green et al., 2010) and empirically assessing the authenticity of ancient DNA sequences (Sawyer et al., 2012) have greatly improved the situation, in practice this concern will never disappear. The importance of controlling for the possibility of contamination is the single most important reason why, to date, convincing ancient DNA research has been dominated by a small number of specialist laboratories with the expertise and facilities to control contamination. All laboratories that carry out this type of work will need to maintain the same level of vigilance currently maintained at specialist laboratories. To reduce contamination at the source, it is also important that archaeologists excavating new samples become trained in handing samples in ways that minimizes contamination. Valuable measures include wearing sterile gloves and a protective suit while excavating remains, not washing the remains, and immediately placing the remains in a sterile plastic bag and refrigerating them prior to shipment to an ancient DNA lab.

A second experimental challenge is the difficulty of identifying human remains that contain preserved DNA. This often requires screening dozens of individuals from multiple sites, with success rates in DNA extraction depending strongly on the conditions experienced by the sample: its age, temperature, humidity, acidity, the part of the body from which it derives, the speed with which it dried after death, whether or not it is from a sample that was rapidly defleshed, and likely other factors that are not currently understood. Modern methods have increased the efficiency of DNA extraction by capturing the short molecules that make up the great majority of any ancient sample (Dabney et al., 2013; Rohland, 2012). In addition, improved library preparation methods have increased the fraction of molecules in an extract that are amenable to sequencing (Fu et al., 2013; Meyer et al., 2012). Nevertheless, it is still the case that there is great variability in success. The secret of successful ancient DNA research is not luck, but hard work and years of experience: screening many carefully chosen and prepared samples until a subset are identified that perform well.

A third experimental challenge is the difficulty of finding a sample that has a sufficiently high proportion of DNA from the bone itself to be economically analyzed. Concretely, the challenge is that for many ancient DNA extracts, the proportion of endogenous DNA is very low, on the order of one percent or less. For example, a recent study on a ~40,000 year-old sample from the Tianyuan site of Northern China had an endogenous DNA proportion of 0.02% (Fu et al., 2013). For such samples, important as they are, it is not economical to simply carry out brute-force sequencing of random genomic fragments and expect to obtain sufficient coverage to make meaningful inferences.

The challenges of ancient DNA research do not end once the wet lab sample preparation is complete, as the datasets that are generated pose formidable computational and analytical challenges.

The first computational challenge is that once useable samples are obtained and sequenced, the data need to be processed. Ancient DNA laboratories, which traditionally have their strongest expertise in archaeology, physical anthropology, or biochemistry, often lack the bioinformatics expertise, data processing power and data storage solutions necessary to handle the millions or even billions of sequences that are generated by modern ancient DNA studies. Moreover, ancient DNA data also requires tailored bioinformatics tools for handling the short sequences that are characteristic of old samples (Kircher, 2012); one cannot simply use existing tools such as SAMtools (Li et al., 2009) or the Genome Analysis Toolkit (DePristo et al., 2011) with the default settings. For example, the sequenced DNA fragments are usually short and degraded, and are expected to have C→T and G→A errors at the ends of the molecules due to cytosine deamination. The data additionally need to be computationally assessed for evidence of contamination, for example by checking the rate of molecules that do not map to the mitochondrial DNA consensus sequence obtained for the sample (Fu et al., 2013). If a sample is determined to be contaminated, the characteristic ancient DNA errors can be leveraged to reduce the level of contamination (assuming that the contamination is not old) by restricting to sequences that contain typical ancient DNA degradations (Meyer et al., 2013; Skoglund et al., 2014).

A second computational challenge for ancient DNA research is that once the data are processed and a sample is determined as likely to be authentic, the data also need to be analyzed using statistical methods that infer population history. While many methods that have been developed to make inferences about population relationships are not biased by the fact that ancient DNA is old and error-prone (Green et al., 2010; Reich et al., 2010; Meyer et al., 2012; Patterson et al., 2012; Prüfer et al., 2014), these methods must be implemented carefully to produce meaningful results. In addition, the methods need to handle complications of ancient DNA, such as the fact that there is rarely deep sequencing data, so that unambiguous determination of genotypes at each position in the genome is often unreliable.

At present, few laboratories have the experimental and computational expertise to address all these challenges simultaneously. As a result, ancient DNA research has been dominated by a few vertically integrated and well-funded laboratories that combine all these skills under the same roof. The high barriers to entry have meant that the full potential of ancient DNA has been largely untapped. In what follows, we sketch out a way forward that we expect will make ancient DNA analysis more accessible to the broad research community, including to archaeologists.

## Democratization of ancient DNA technology

The usual paradigm in ancient DNA analysis of the nuclear genome has been to identify a sample that has a high enough proportion of DNA to deeply sequence. Such golden samples are rare, and are typically identified only after laborious screening of many dozens of samples that have low proportions of human DNA. Once a sample is found that has an appreciable proportion of human DNA, it is typically sequenced to as high a coverage as possible (sometimes the limited number of starting molecules in many ancient DNA libraries is the main factor limiting sequencing depth). All of this is very expensive, and has been an important barrier to entry into ancient DNA work for less well-funded laboratories.

Whole genome sequences, however, are not required for most historical inferences. In a paper that was important not only for what it showed about history but also for what it showed about the quantity of information that can be extracted from small amounts of data, Skoglund et al., (2012) showed that even low levels of genome coverage per sample – 1–5% of genomic bases covered in their case – was sufficient to support profound historical inferences.

Following the example of Skoglund et al. (2012), we propose that a way to make ancient DNA analysis accessible to a much larger number of laboratories is to use targeted capture approaches that enrich a sample for human DNA. Enrichment strategies have already been shown to be extremely effective for ancient DNA analysis. In one approach to target enrichment, Carpenter et al., (2013) showed that it is possible to take RNA baits obtained by transcribing a whole human genome, and hybridize them in solution to an ancient DNA library, thus increasing the fraction of human DNA to be sequenced by several orders of magnitude. Fu et al., (2013) showed that it is possible to synthesize oligonucleotide DNA baits targeted at a specified subset of the genome—the entire mitochondrial DNA sequence, or the coding sequences of all genes—and to hybridize to ancient DNA libraries in solution to enrich a library for molecules from the targeted subset of the genome. This strategy has been used to obtain a high quality mitochondrial genome sequence from a ~400,000 year-old archaic human (Meyer et al., 2013), as well as approximately 2-fold redundant coverage of chromosome 21 from the ~40,000 year old Tianyuan sample from northern China (Fu et al., 2013).

We are particularly enthusiastic about the possibility of adapting a technology like that of Fu et al., (2013) to enrich human samples for panels of several hundred thousand single nucleotide polymorphisms (SNPs) that have already been genotyped on large panels of present-day samples. This is a sufficient number of SNPs that it would allow for high-resolution analysis of how an ancient sample relates to present-day as well as other ancient samples. The strategy has two major advantages. First, through enrichment, it allows analysis of samples with much less than 10% human DNA that are not economical for whole genome sequencing studies. Second, it requires about two orders of magnitude less sequencing per sample to saturate all its targets (Table 1). We caution that there are some questions—for example, estimation of population divergence times—that rely on identification of sample-specific mutations and may be better addressed with whole genome sequencing than by a SNP capture experiment. Nevertheless, we believe that a capture experiment can answer the substantial majority of questions related to population history or natural selection that are addressable with genetic data, while allowing larger numbers of samples to be analyzed for the same cost.

**Table 1.** Comparison of two strategies for ancient DNA studies of history

|  | Whole genome sequencing | Capture and sequencing of 300,000 SNPs |
| --- | --- | --- |
| Number of samples that need to be screened* | 1 | 1/10 |
| Amount of sequencing that needs to be performed* | 1 | 1/100 |
| High resolution inference of pop. relationships | Yes | Yes |
| Allows studies of selection | Yes | Yes |
| Variants specific to sample can be discovered | Yes | No |
| Works for samples with low % of human DNA | No | Yes |

\* These are rough qualitative estimates based on discussion with colleagues. The estimated amount of work required for capture 300,000 SNPs is expressed as a fraction of that required for whole-genome sequencing.

An important goal for the coming years should be to make DNA a tool that will become fully accessible not just to smaller labs but also to archaeologists. In this regard, it is important to encourage interaction and collaboration between the genetic and archaeologist communities to develop standards for the interpretation and use of genetic data in answering questions relevant to archaeology.

Members of the archaeologist community are already sophisticated users of other scientific technologies for analyzing ancient biological remains such as carbon 14 analysis (for date estimation) and stable isotope analysis (for making inferences about diet). Currently, when archaeologists have a sample that they wish to analyze, they send it to specialist (sometimes commercial) laboratories, which then provide a carbon date and/or an isotope analysis, along with a report giving interpretation. We envision a future in which ancient DNA analysis services could be provided in a similar way. Concretely, archaeologists would be able to send skeletal material to a specialist laboratory for analysis, where it can be tested for ancient DNA using up-to-date protocols. If ancient DNA is detected, a whole mitochondrial genome sequence can be produced and a sex can be determined. For ancient samples that produce uncontaminated DNA, whole genome data can be produced.

To make such a future possible, it will be necessary to build up a database of present-day human and ancient DNA samples all genotyped at the same set of SNPs to which any new sample can be compared. In addition, it will be necessary to write software to automatically compare a new sample to samples in the database (Alexander et al., 2009; Patterson et al., 2012; Price et al., 2006; Raj et al., 2013), which will be useful for producing a report for archaeologists on how the sample relates to diverse present-day humans, and to other ancient samples. Because of the subtleties of interpreting genetic data, it will important be necessary for archaeologists to work collaboratively with population geneticists in interpreting the results of such studies. Such a report would, however, be a useful tool for giving archaeologists a first impression of the biological ancestry of their samples.

We particularly wish to highlight the potential of ancient DNA as a direct tool for archaeological analysis, which will be useful for elucidating the population structure and patterns of relatedness within a particular archaeological site. If many individuals from an archaeological culture can be analyzed in a way that correlates the genetic information—sex, patterns of relatedness, and ethnic affiliation—to archaeological information derived from grave goods, status symbols, and other objects, it seems likely to provide new insights. A particular area of opportunity is the identification of outlier individuals. Genetic data should make it easy to detect individuals that are outliers in terms of their ancestry relative to others at nearby sites, which could be informative about migration rates and sex biases in these rates.

## Conclusions

We have argued that it will likely to be fruitful to re-examine many aspects of human population history and the history of natural selection from a perspective in which population movements play a central role. We have further argued that advances in ancient DNA are likely to have an important impact on how we think about using genetic data to address questions about human history.

We conclude by presenting three questions that we believe can be answered in the coming years by genetic studies of human history and variation; this list is not comprehensive, and many other important questions surely can also be addressed:

1. What is the genetic legacy of the inhabitants of eastern Africa before the arrival of Bantu speakers? The population genetic structure of sub-Saharan Africa has been transformed by the expansion of Bantu-speaking agriculturalists over the last 3,000 years. As a concrete example, the population structure of eastern Africa prior to this expansion remains controversial (Morris, 2003; Schepartz, 1988). Ancient DNA from samples prior to the Bantu expansion could settle this debate. While the climate in much, but not all, of sub-Saharan Africa is less favorable for preservation of DNA than that of northern Eurasia, it is not clear whether enough African samples have been tested to determine whether or not ancient DNA analysis works. Moreover, the time scale of the Bantu expansion is only several thousand years rather than tens of thousands of years. In light of continuing technological progress in ancient DNA extraction (including the successful extraction of DNA from a ~400,000 year old sample in Spain (Meyer et al., 2013)), we are hopeful that ancient DNA studies of some African samples within the last few thousand years may become possible.
2. When were the geographic patterns at loci under positive selection established? Loci under positive selection in humans have allele frequencies with characteristic geographic patterns, which have been interpreted as “west Eurasian”, “east Asian” and “non-African” selective sweeps (Coop et al., 2009). Were these patterns established tens of thousands of years ago during the initial human peopling of the globe by anatomically modern humans? Or were they established after later population movements? Resolving these questions will become possible once substantial numbers of ancient samples are genotyped. Work in this area is only just beginning (Wilde et al., 2014).
3. Was the spread of Indo-European languages caused by large-scale migrations of people or did language shifts occur without extensive population replacement? For times prior to the invention of writing, it will never be possible to directly relate the language that people spoke to their remains. Nevertheless, genetic data may still informative about the historical events that accompany linguistic expansions. The origins of the Indo-European languages—spoken across Europe and West, Central, and South Asia—is a particularly important question (Anthony, 2007; Mallory, 1997). These languages likely spread across Eurasia and diversified within the last 5-10 thousand years. Was this spread of languages caused by a large-scale movement of people? It now may be possible to obtain ancient DNA from cultures known to have spoken Indo-European languages—for example, Hittites and Tocharians—and to compare these populations with their neighbors to determine whether they harbor a genetic signature specific to the Indo-European speakers. It may also be possible to search for the spread of (currently hypothetical) Indo-European genetic signatures in Europe and India at the times when various hypotheses have suggested that they may have arrived.

In conclusion, genetic studies of the human past are at the beginning of an era in which genome-wide studies of modern and ancient DNA samples will make it possible to ask and answer questions about human history and natural selection that could not be addressed before. We expect that these approaches will provide new insights into important and still controversial issues.

## Acknowledgements

We thank Henry Harpending and Alan Rogers for a helpful discussion about the origins of the serial founder effect model. We are grateful for discussions and comments on the manuscript from David Anthony, Joachim Burger, Qiaomei Fu, Alexander Kim, Iosif Lazaridis, Iain Mathieson, Nick Patterson, Molly Przeworski, Nadin Rohland, Pontus Skoglund, Johannes Krause, Richard Meadow and Wolfgang Haak. DR was supported by NSF HOMINID grant BCS-1032255 and NIH grant GM100233 and is an Investigator of the Howard Hughes Medical Institute.

